# Habitat Protection Indexes - new monitoring measures for the conservation of coastal and marine habitats

**DOI:** 10.1101/2021.06.09.447318

**Authors:** Joy A. Kumagai, Fabio Favoretto, Sara Pruckner, Alex D. Rogers, Lauren V. Weatherdon, Octavio Aburto-Oropeza, Aidin Niamir

**Affiliations:** Senckenberg Biodiversity and Climate Research Center, Frankfurt am Main, Germany; Centro para la Biodiversidad Marina y Conservación, A.C., La Paz, Baja California Sur, Mexico; Universidad Autónoma de Baja California Sur, La Paz, Baja California Sur, Mexico; UN Environment Programme World Conservation Monitoring Centre (UNEP-WCMC), Cambridge, United Kingdom; REV Ocean, Lysaker, Norway; Scripps Institution of Oceanography, University of California San Diego, La Jolla, California, USA

## Abstract

A worldwide call to implement habitat protection aims to halt biodiversity loss. To monitor the extent of coastal and marine habitats within protected areas (PAs) in a standardized, open source, and reproducible way, we constructed the Local and the Global Habitat Protection Indexes (LHPI and GHPI, respectively). The LHPI pinpoints the jurisdictions with the greatest opportunity to expand their own PAs, while the GHPI showcases which jurisdictions contribute the most in area to the protection of these habitats globally. Jurisdictions were evaluated to understand which have the highest opportunity to contribute globally to the protection of habitats by meeting a target of 30% coverage of PAs with Areas Beyond National Jurisdiction (ABNJ) having the greatest opportunity to do so. While we focus on marine and coastal habitats, our workflow can be extended to terrestrial and freshwater habitats. These indexes are useful to monitor aspects of Sustainable Development Goal 14 and the emerging post-2020 Global Biodiversity Framework, to understand the current status of international cooperation on coastal and marine habitats conservation.

## Introduction

Marine habitats face simultaneous global and local pressures with more than half of the ocean experiencing significantly increasing cumulative impacts in the last decade^1^. These habitats provide critical services to humans (i.e., nature’s contributions to people) spanning regulating (e.g., coastal protection and CO2 sequestration), provisioning (e.g., food), supporting (e.g., habitat provision), and cultural services (e.g., science and well-being). Yet habitat destruction has continued into the 21st century, with the loss of one-third to one-half of vulnerable coastal and marine habitats globally, accompanied by a reduction in nature’s contributions to people^2^. In order to solve this worldwide crisis, there are calls from various organizations including the High Level Panel for a Sustainable Ocean Economy3 and the Intergovernmental Oceanographic Commission (IOC) through the UN Decade of Ocean Science for Sustainable Development^4^ for the use of data to drive decision-making. However, there is no standardized database or indicator available that reports on how much coastal and marine habitats fall under protected areas (PAs) or other effective area-based conservation measures (OECMs) within the jurisdiction of each country or territory.

PAs and OECMs^5^ are critical to maintain the health of ecosystems and their contributions to people. When MPAs are properly planned, designated, implemented, and managed6,7, habitat integrity is secured, depleted fish populations can rebound and biodiversity can increase, which can lead to spillover effects^8,9^. Regulation and correct placement of MPAs are fundamental to protecting ecosystems effectively. Although undesirable outcomes can result from ineffective implementation of MPAs^10–12^, it has been shown that 90% of the maximum potential biodiversity benefits from MPAs can be achieved by strategically protecting 21% of the ocean (43% of EEZs and 6% of the high seas)^13^.

As of May 2021, 7.74% of the global ocean area was covered by MPAs^14,15^ and 2.7% was within fully protected areas^16^. Target 2 of the zero draft of the post-2020 global biodiversity framework currently under negotiation proposes that by 2030 at least 30% of the earth’s surface should be protected or conserved through well connected and effective system of protected areas and other effective area-based conservation measures with the focus on areas particularly important for biodiversity17,18. Whether or not this is agreed upon, setting targets for ecosystem conservation is essential for policy-makers19. For example, it has been demonstrated that by prioritizing food provisioning by the ocean, the full protection of 28% of the ocean will provide a net gain of 5.9 MMT of seafood and incidentally secure 35% of biodiversity and 27% of potential carbon sequestration13.

Only 11% of PAs, representing 18.29% of global protected area coverage, have been assessed for their management effectiveness and only 15.4% of countries meet their targets of assessing at least 60% of PAs for their effectiveness^15^. Transparency in protection efforts and effectiveness must be improved so policymakers can grasp the current conditions, possible scenarios, and make informed decisions to meet international policy commitments20. Further, to deliver the maximum potential biodiversity benefits, PAs and OECMs need to include a sufficient representation of the world’s habitats and species15. Therefore, we provide a comprehensive dataset and open-source workflow to standardize the amount of marine and coastal habitats within PAs or OECMs at the local and global level, monitoring the necessary representation of diverse habitats within PAs.

We define two new indexes: the global habitat protection index (GHPI) and the local habitat protection index (LHPI). The GHPI is the area of habitats within PAs or OECMs in a jurisdiction divided by the global extent. The LHPI is defined as the average area of habitats within PAs or OECMs in a jurisdiction divided by the extent of the habitats in the same jurisdiction. The indexes can be calculated at any resolution larger than a few kilometers, without the need of proprietary software, as long as the underlying data have a higher resolution than what is used. While we focus on marine and coastal habitats, any habitat layer can be used in our workflow. Our approach applies equally to areas beyond national jurisdiction and could be extended to terrestrial and freshwater habitats. The indexes shed light into the effort governments are putting towards monitoring marine and coastal habitat conservation and can be used for both global level policy and national statistical offices.

## Results

On a global level, more than 42% of warm-water corals, mangroves, and saltmarshes fall under PAs or OECMs, while 27– 29% of seagrasses and cold-water corals are under PAs or OECMs (Figure 1). Knolls and seamounts have the least area covered by PAs or OECMs, with a total of 8%. If we consider just the ABNJ, where 67% of the extent of knolls and seamounts are located (Figure 1a), only 0.8% are under PAs or OECMs. The discrepancy between protection efforts in coastal regions versus ABNJ is a result of an overall protection focus on national waters and of the underlying challenges to create international conservation efforts within ABNJs. Since numerically half of the selected habitats are found within ABNJ (cold corals, seagrasses, knolls, and seamounts) it is important to implement new measures to protect these areas.

**Figure 1:**
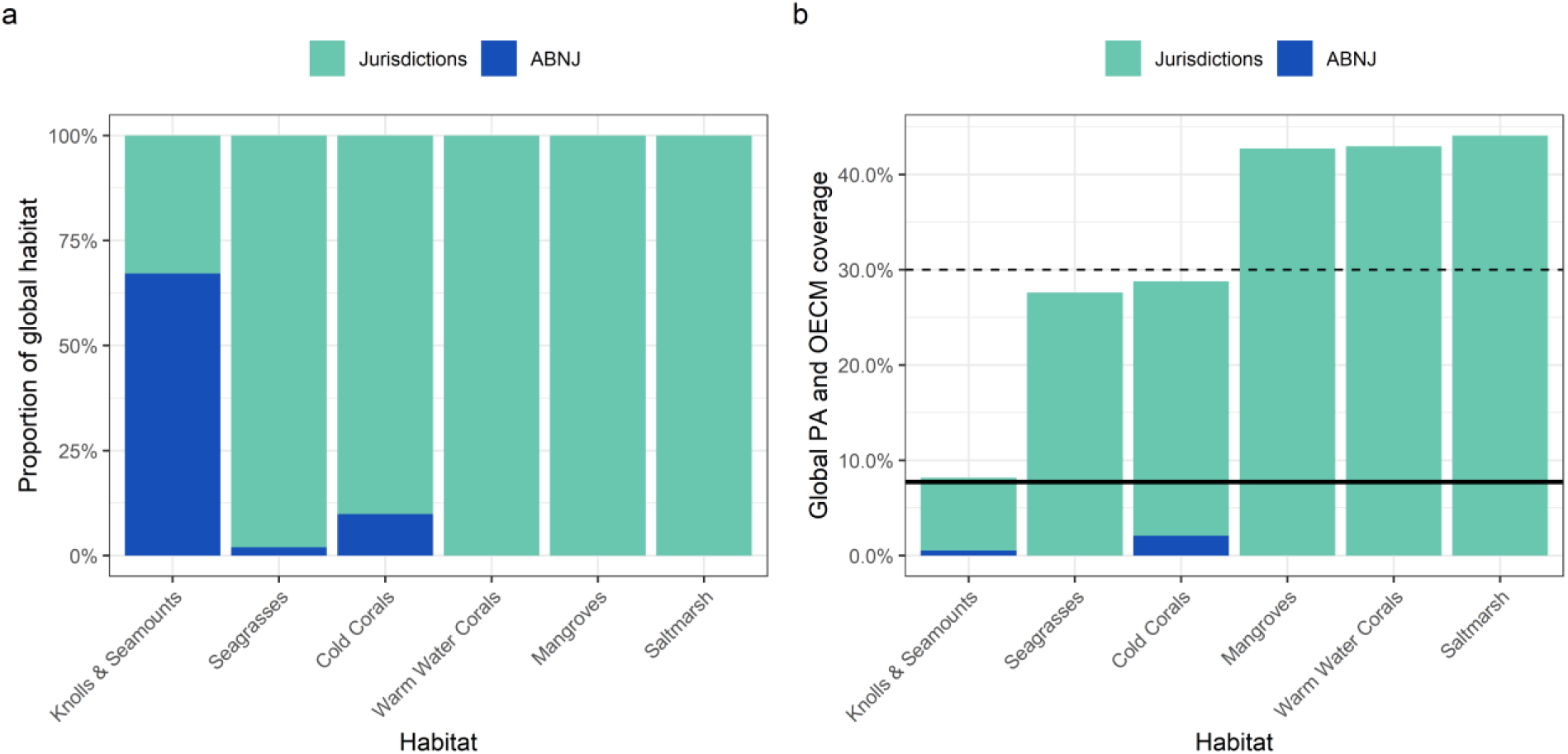
Overview of the distribution and protection of the six habitats considered over the jurisdictions and ABNJ. (a) Illustrates the extent distribution of each habitat between jurisdictions and ABNJ. Warm water corals, mangroves, and saltmarsh do not occur in ABNJ; (b) Represents the global coverage of PAs and OECMs for each habitat divided between jurisdictions and ABNJ. The dashed line at 30% represents the target that 30% of the ocean should be conserved by 2030, while the solid black line at 7.74% represents the current protection of the ocean surface area according to UNEP-WCMC^15^.

At the jurisdiction level, we selected 242 jurisdictions that include at least one of the habitats used in the analysis. The GHPI measures how much a jurisdiction contributes to the total extent of the marine and coastal habitats we consider within PAs or OECMs globally. The top five jurisdictions with the highest average GHPI value are Australia, Canada, Mexico, Spain, and Indonesia (Figure 2), which are within the top 10 jurisdictions with the largest EEZ area except for Spain. Notably, the GHPI has a highly right-skewed distribution, meaning that just a few jurisdictions contribute extensively to habitat protection on a global scale. These top five jurisdictions together contribute 15.8% (GHPI score of 0.158) of the average global protection of the considered marine and coastal habitats by extent.

**Figure 2:**
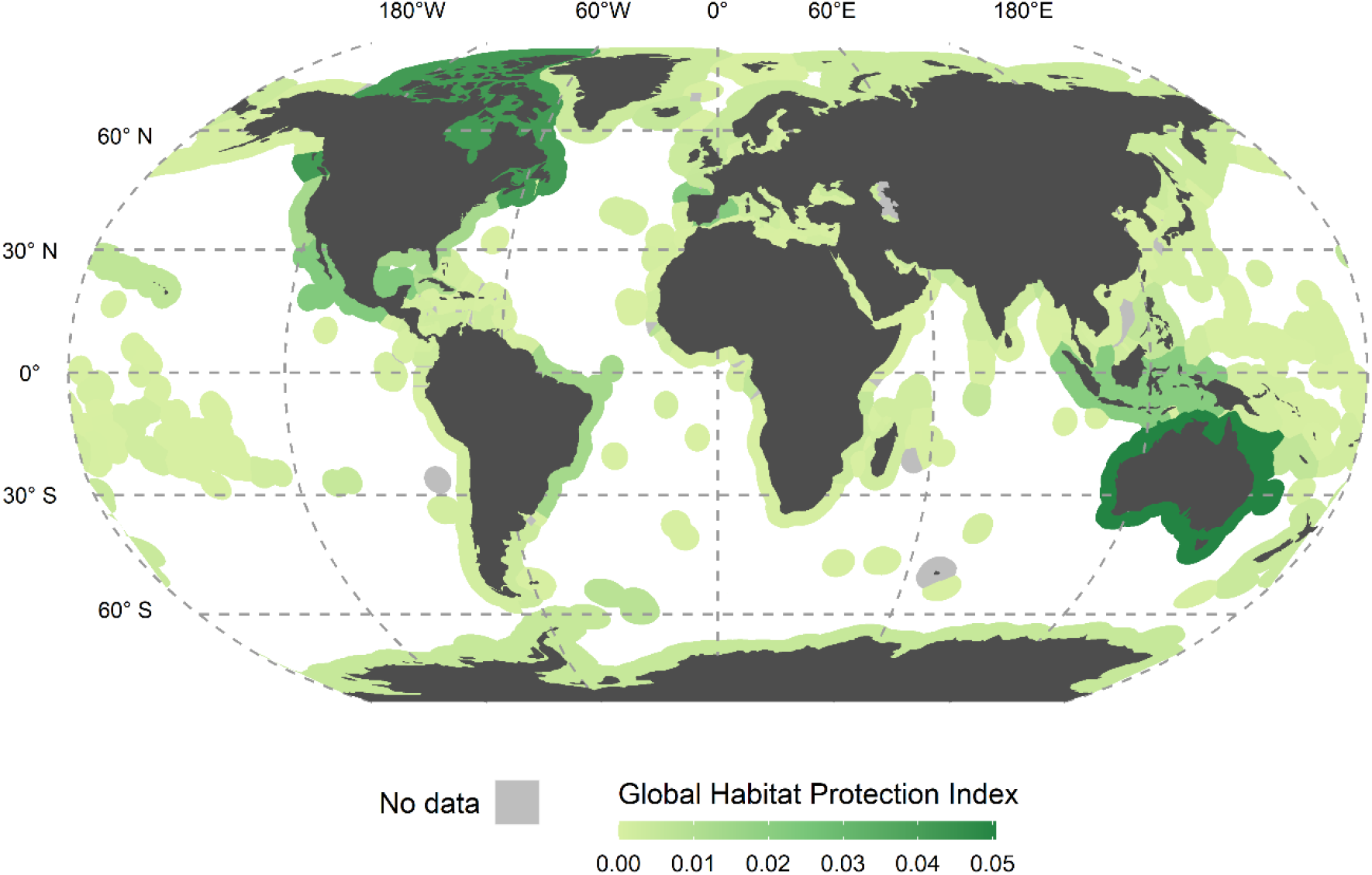
The Global Habitat Protection Index (GHPI). GHPI illustrates the contribution of jurisdictions to the global coverage of six marine and coastal habitats by PAs or OECMs, ranging from yellow-green (low contribution) to dark green (high contribution). The GHPI is calculated by taking the average of each habitat specific GHPI presented in the Supplementary Information 2, habitats that do not occur in the jurisdiction’s extent are not included in the calculation. The distribution of the index is highly right-skewed, with a few jurisdictions with comparatively very high scores and many jurisdictions with low scores. The ABNJ index value is not depicted for clarity. Index ranges from 0 to 1, but only 0 to 0.05 is depicted here due to no jurisdictions scoring higher than 0.05.

Comparatively, the LHPI, which measures on average how much coastal and marine habitat in a jurisdiction is under a PA or OECM, also has a right-skewed distribution. The LHPI varies greatly over the jurisdictions (Figure 3) and also for the six selected marine and coastal habitats when calculated for each habitat individually (Supplementary Information 2). More than half of jurisdictions have an LHPI value of less than 0.27 and 25% of jurisdictions (61) have an LHPI value higher than 0.50. Nine jurisdictions have an LHPI value of 1, indicating that all their habitats occur completely within PAs, while 30 jurisdictions have a value of 0, where none of these habitats are within PAs. Many of the top LHPI values are held by island jurisdictions. For example, Bouvet Island, Johnston Atoll, and Saint Barthélemy all have a LHPI value of 1.

**Figure 3:**
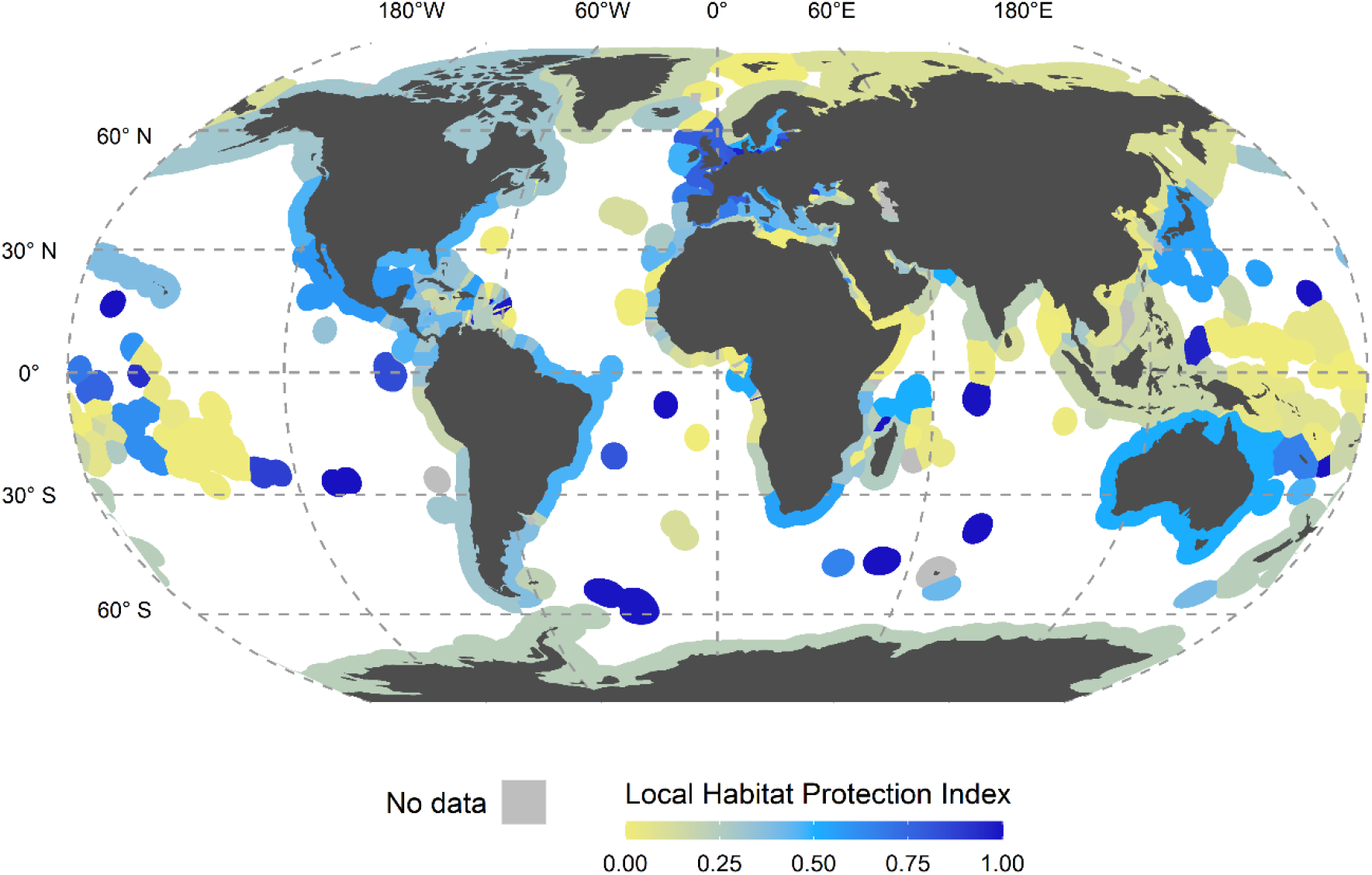
The Local Habitat Protection Index (LHPI). LHPI illustrates how much a jurisdiction is covering the six marine and coastal habitats considered with PAs or OECMs compared to the maximum habitat extent, ranging from yellow (low contribution) to dark blue (high contribution). The index ranges from 0 to 1, indicating no to 100% coverage of PAs or OECMs. The LHPI is calculated by taking the average of each habitat specific LHPI presented in the Supplementary Information 2. Habitats that do not occur in the jurisdiction’s extent are not included in the calculation. The ABNJ index value is not depicted for clarity.

Finally, an analysis was conducted to calculate an additional benchmark called the targeted GHPI, which measures how distant jurisdictions are to protecting 30% of their habitats (Figure 4; Supplementary Information 3 for habitat specific figures). The targeted analysis of the GHPI subtracts 30% of each habitat from the GHPI (see methods), so that jurisdictions with positive values on average have more than 30% of the selected habitats within PAs, whereas jurisdictions with negative values are not meeting this goal. The top five jurisdictions with positive targeted GHPI values are Australia, Canada, Spain, Mexico, and Brazil. Interestingly, the ABNJ rank the lowest in this analysis even though they rank 10th highest on the GHPI. This is because three of the six habitats considered fall within the high seas, and they have a large area including cold-water corals, and knolls and seamount habitats. However, only a small proportion of the high seas are within protected areas, leading to this disconnect. The index values for the LHPI and GHPI, and the results of the targeted analysis of the GHPI calculation can be found in the accompanying repository.

**Figure 4:**
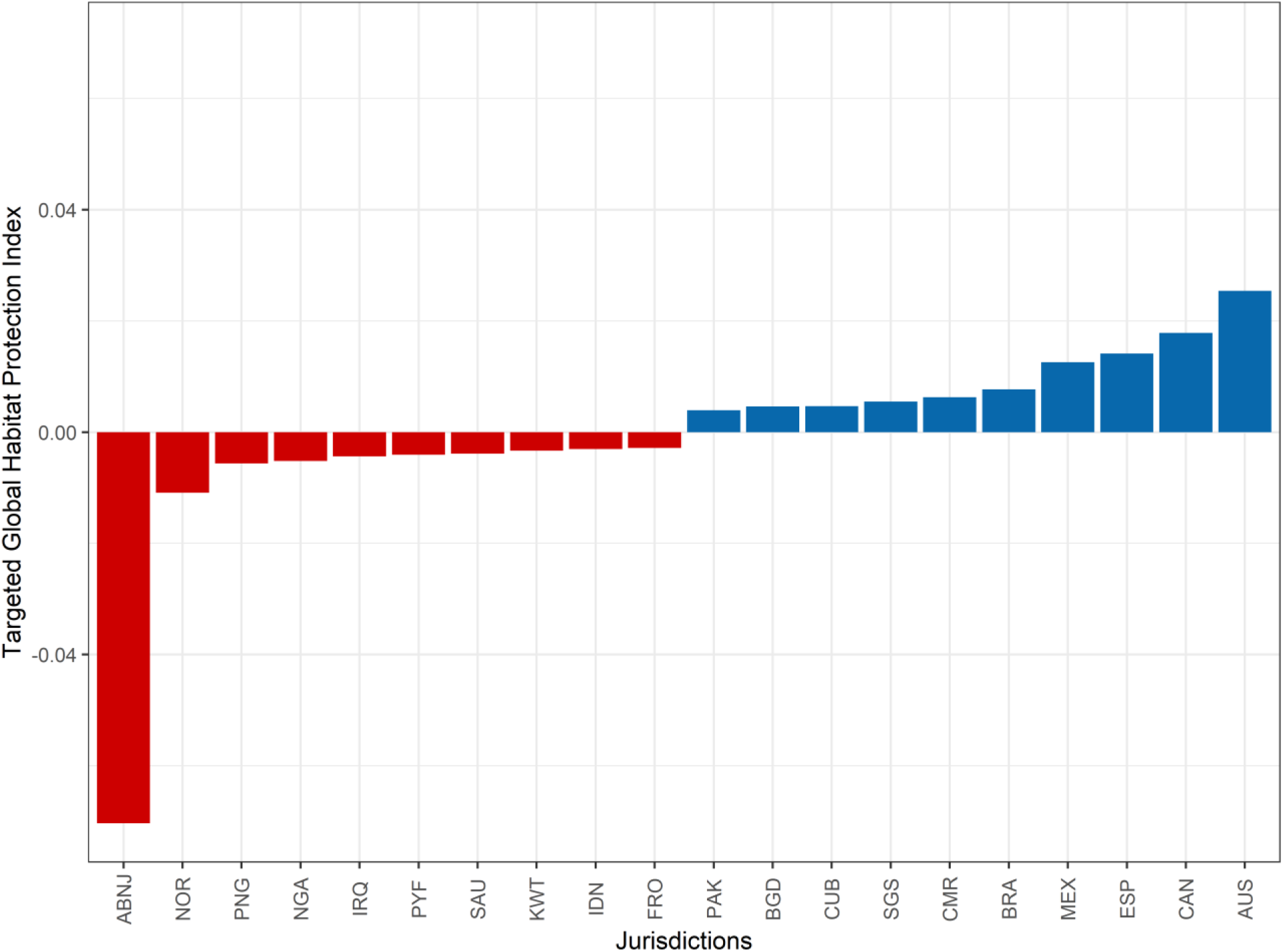
The top 10 and bottom 10 out of 242 jurisdictions ranked according to the results from the targeted analysis of the GHPI illustrating whether these countries have on average 30% of their habitats within PAs or OECMs. Jurisdictions in blue (above 0) have on average more than 30% their habitats extent within PAs or OECMs, while jurisdictions in red (below 0) have on average less than 30% of their habitats’ extent within PAs or OECMs. Jurisdiction names correspond to ISO3 names except for the ABNJ.

## Discussion

There are 23 international conventions that relate to the protection of the marine environment and biodiversity with 5 of these requiring the implementation of MPAs21. Setting of targets for effective protection of marine habitats which both protect nature and secure nature’s contributions to people are increasingly seen as critical in ensuring progress toward meeting treaty commitments. Aichi Target 11 and SDG Target 14.5 set targets to conserve at least 10% of marine and coastal areas by 2020 reflecting a shift to more target-driven conservation policy at international level although still debated. Warm-water corals, mangroves, and saltmarsh all have more than 30% of their extent within PAs or OECMs with seagrasses and cold corals approaching 30%, which is a great accomplishment and reveals dedicated effort to their conservation. However, the protection of the total global ocean area is still at 7.74%, with only 1.18% of the ABNJ covered by PAs or OECMs, falling short of the 10% Aichi goal 11 previously set for 202015.

To plan how to achieve such targets and monitor progress, useful tools such as data platforms are required to assess progress in biodiversity conservation, such as the Protected Planet initiative, in which policymakers and managers can have confidence enabling effective collaboration. Transparent tools which process data within these platforms will help standardize statistics and information. Such tools have been lacking until recently, but the habitat protection indexes presented here provide one of the first steps to fill this gap. Combined with systematic spatial conservation planning that allows policy to direct a balance of tradeoffs for other priorities, such as climate mitigation and resource extraction (e.g., ^13^), and consideration of human rights of local communities and indigenous people in the decision-making process22, the LHPI and GHPI present a consistent way of measuring progress in establishing PAs that meet the requirement to enhance coastal and marine habitat protection. The workflow to calculate the indexes presented here is highly adaptable and can include a much wider range of habitats including terrestrial and freshwater ecosystems, as well as other area-based conservation targets. Additionally, the inclusion of ABNJ in the indexes is extremely important given current discussions on a new implementing agreement for the United Nations Convention on the Law of the Sea to protect marine biodiversity in areas beyond national jurisdiction and thus the whole ocean^23^.

Jurisdictions have direct control over their LHPI, as increasing the protection of their marine and coastal habitats will directly increase the index score. Small countries and territories with limited area may see large improvements in their LHPI through a few additional PAs, while their GHPI score will not increase much from this effort. Within the targeted GHPI analysis, any jurisdiction which decides to protect more than 30% of their habitat extent can move from a negative to a positive score. For jurisdictions with large extents of habitats such as Australia, more effort is required to increase their LHPI, but small improvements in their LHPI will correspond to significant increases in their GHPI score, contributing greatly to the global efforts of protecting marine and coastal habitats. It is important to note that a human rights-based approach is absolutely essential in meeting the targets set and improving the index scores.

Considering the targeted analysis of the GHPI (Figure 4), the jurisdictions that rank lowest in the analysis (High Seas, Norway, Papua New Guinea, Nigeria, and Iraq) represent a great opportunity to further expand PAs to 30% coverage of marine habitats within their territorial waters and coast. The jurisdictions that score highest have the opportunity to improve the effectiveness of their PAs to adequately protect these marine habitats and reduce surrounding pressures, especially as they contribute significantly to the total global extent of these habitats.

The analysis presented here is sensitive to the choice of coastal and marine habitats that are included in the indexes. We selected these 6 habitats based on the availability of high-quality spatially explicit global data recognized by the scientific community. Each of the marine habitat datasets are from peer-reviewed publications that were validated individually by the UN Environment Programme World Conservation Monitoring Centre (UNEP-WCMC) following their data standards. Over time, the workflow will be updated and improved yearly to strengthen data coverage, and if additional high-quality data on habitats that can be used for policymaking emerge these will be included, thus ensuring the indexes stay up to date and relevant. The original analysis with the same habitats will also be repeated to ensure that a consistent time series of the indexes values is provided.

An important consideration when using these indexes is that habitat extent that spatially aligns with a protected area or OECM does not necessarily mean that a particular habitat is protected. For example, there are PAs that exclude fishing and cover some coastal extent with mangroves but do not prevent mangrove deforestation by other activities. Additionally, because of the buffering of points within the workflow, some of the habitat that is counted as protected may fall near a PA or OECM but not within it. Nevertheless, in our analysis it is assumed that habitats that fall within a protected area or OCEM will be better conserved than habitats not within a protected area, as the primary purpose of PAs is conservation, OECMs provide conservation benefit, and these areas often are sustainably managed by local communities and indigenous people who live on them^24,25^.

In summary, the LHPI and GHPI indexes report detailed information for policy makers, the scientific community, and stakeholders to understand the state of protection of marine and coastal habitats at both the global and local level. Based on open-source programming and datasets, the reproducible and scalable workflow has been developed to allow for others to calculate the indexes for any areas or habitats of interest and repeat our analysis for any target. The indexes will be updated annually, to ensure continued relevance and the provision of a time series to track how the world is progressing towards the goals defined by global policy, such as aspects of the Sustainable Development Goal 14, therefore bringing to the forefront the importance and status of conserving critical marine and coastal habitats.

## Methods

Using the free and open-source programming language, R, and open-access datasets frequently used by the scientific community, two indexes were created focused on the protection of six marine and coastal habitats, and paired with an analysis considering the proposed target of 30% protection. To calculate the indexes, we have created a standardized and reproducible workflow described below following the FAIR data management principles. All datasets included in the workflow are listed in Supplementary Information 1 and the code for the workflow can be accessed in the Zenodo repository (https://doi.org/10.5281/zenodo.4694821).

### Data Collection and Processing

To calculate the coverage of PA and OECMs for each of the habitats, the first step of the workflow was to download, clean and filter the Protected Planet Initiative dataset of protected areas14 and the World Database on Other Effective Area-based Conservation Measures (WD-OECM)26. We followed the Protected Planet Initiative method for the PAs described on https://www.protectedplanet.net/en/resources/calculating-protected-area-coverage but adapted it to the R programming language. We removed UNESCO Man and the Biosphere (MAB) Reserves, PAs that were reported with a status of ‘Proposed’ or ‘Not Reported’, and PAs reported as points with no reported area. Points with a reported area were buffered to create polygons matching the reported area for both PAs and OECMs. Next, the PAs and OECMs were re-projected to the Behrmann equal-area projection. All protected area layers were then converted into a raster of 1 km2 pixels and merged together.

To develop our workflow, we selected 6 coastal and marine habitats from an authoritative source of internationally recognized data, the UNEP-WCMC. We selected these 6 habitats based on their ecological importance and availability of high-quality spatially explicit global data recognized by the scientific community. The Ocean Data Viewer supplies downloadable data after an approved data standard procedure prior to any dissemination. The six habitats selected are cold-water corals, warm-water corals, knolls and seamounts, mangroves, saltmarshes, and seagrasses and were all downloaded from the Ocean Data Viewer. The union of knolls and seamounts’ base area was used. Habitat layers are projected and converted into rasters. For points, we buffer them to create polygons with an area equaling the reported area; if the points had no area assigned, we assumed they have a 1 km2 extent.

The habitat rasters were then intersected with the PAs and OECMs raster. Therefore, for each habitat, two layers were produced. One representing the habitat in 1 km2 pixels, and a second representing the habitat within PAs or OECMs. We then extracted the number of pixels within each area of interest using parallel processing for all of the resulting layers. Our areas of interest were the union of countries or territories and their respective EEZs available online at https://www.marineregions.org/.

We used the union of land and their EEZs remaining at the territory level to ensure that the extent of coastal habitats such as mangroves and saltmarsh were not clipped by the coastline. Landlocked countries, as well as disputed and joint-regime areas were also filtered from the dataset, therefore the global extent used in the indexes’ calculation is the sum of the habitat extent of the remaining jurisdictions. The global statistics reported in Figure 1, were calculated based on the number of pixels within each habitat layer, independent of the areas of interest. We also calculated the number of pixels within ABNJ with the same method. While the process of rasterization of polygon data may decrease area accuracy, due to the very high-resolution grid used, 1 km2, very little accuracy was lost. Comparing the current area reported for each of these habitats available from Ocean+27, we had less than a 0.5% difference of area from this conversion.

No-take areas and effectively managed areas can easily be removed or included in our workflow. Considering that numerically 99.06% of the PAs in the World Database on Protected Areas (WDPA) are assigned to the categories “not applicable” or “not reported” regarding their no-take status, it was decided to not include any data on no-take areas for our indexes. For example, in Mexico, their national databases have 329,875 km2 of area within no-take PAs (4.8% of the EEZ)^28,29^, yet zero no-take area is reported in the WDPA in Mexico. Once there is more comprehensive global data on no-take statuses, these areas can be easily included as a separate category of our indexes because of its demonstrated conservation importance. Other effective area-based conservation measures (OECMs) are included in the HPI calculation as they are explicitly mentioned in Target 2 of the post 2020 biodiversity framework currently being negotiated17,18. The current World Database on Other Effective Area-based Conservation Measures (WD-OECM)26 only has data reported from five countries and territories and 506 records, but further efforts to collect data on OECMs having initiated following the 14th Conference of Parties of the Convention on Biological Diversity in 2018 (Decision 14/8)30.

### Index Calculations and Considerations

Using the resulting data, we first calculated the average percent of the marine and coastal habitats falling within a PA or OECM for a jurisdiction. First, we divided the total number of a specific habitat’s pixels within PAs or OECMs by the total number of the same habitat’s pixels for each area of interest (Supplementary Information 1, Figures 1b-6b). We then average the resulting numbers for the six habitats selected, mangroves, seagrasses, saltmarsh, cold corals, warm-water corals and knolls and seamounts, resulting in the Local Habitat Protection Index (LHPI, Figure 3). The LHPI index measures how much a jurisdiction is covering their habitat extent with PAs or OECMs compared to their maximum amount. The next index, the global habitat protection index (GHPI), measures how much a jurisdiction is contributing to the total extent of habitats within PAs or OECMs globally. The GHPI is calculated by dividing the extent of each habitat within PAs or OECMs within a jurisdiction by the global extent of that habitat (sum of each jurisdiction’s habitat extent) (Supplementary Information 1, Figures 1a-6a). We then average this value for the six habitats (Figure 4). The theoretical range for both indexes is from 0 to 1.

Finally, we argue that at least 30% of each of these habitats should be protected as a result of the high value of their contributions to people and high importance for biodiversity protection coupled with the current discussions of the post-2020 global biodiversity framework17. With this in mind, we created an analysis which evaluates how well jurisdictions are meeting this goal. Each jurisdiction’s target extent of habitat within PAs or OECMs is calculated by multiplying its global fraction of habitat by 0.3 and subtracting this from the GHPI (habitat-specific). This calculation is done for each habitat (Supplementary Information 3, Figures 7-12) and the average of all habitats are presented in Figure 4.

The workflow, on which the indexes are based, runs on a 1 km2 scale, thus in the rasterization process, small patches of habitat polygons that do not intersect the centroid of each raster cell will not be included. If the habitat polygon does intersect with the centroid it will be rasterized to a 1 km2 resolution. Additionally, the final indexes are the average of the protection for all habitats present in the jurisdiction being considered. Because of this, some areas or countries with very little habitat in small patches have a value of 1 for the LHPI because the limited areas of the habitat that were rasterized fall within a PA or OECM. For example, Bouvet Island has 50 pixels of knolls and seamounts in the dataset and all fall within a protected area, additionally no other habitats are present according to the datasets, therefore the jurisdiction has an LHPI value of 1. Because of this rasterization process, there may be small patches of habitat that are not under a protected area within the country or territory but are not reported here.

## Data Availability

The final dataset in CSV format with the indexes is available on Zenodo (https://doi.org/10.5281/zenodo.4694821), consisting of the LHPI and GHPI, and the results of the targeted analysis of the GHPI, for each jurisdiction. An additional dataset with more detailed information is included where the indexes for each habitat per jurisdiction is reported alongside the habitat specific 30% analysis results. These indexes will be updated annually and include trends over time after a few years. Additionally, these indexes will be further improved overtime with additional habitats as more detailed spatial habitat data is openly available on a global scale, and the protected areas database continues to be updated with higher resolution data and the statuses of no-take PAs.

## Code Availability

The workflow consisting of several R scripts used to produce these datasets can be found in the Zenodo repository together with the data files and supplementary figures. The ongoing development of code can be tracked on Github here: https://github.com/jkumagai96/Marine_Habitat_protection. All the data presented in the figures in the manuscript can be reproduced with the available code which was originally drafted and run in R Studio (version 1.4.1103 – “Wax Begonia” Windows) with R version 4.0.3. The packages used are managed through *renv* with these associated files are available on Github. Please note that the workflow requires the user to download all the habitat data and the union of EEZ and country polygons from the data sources listed in Supplementary Information 1 (Supplementary Information 1 Table 1).

## Acknowledgements

This work represents an extension of the analysis presented first in the 10th Blue Paper, “Critical Habitats and Biodiversity: Inventory, Thresholds and Governance,” for the High-Level Panel for a Sustainable Ocean Economy. We thank Jessica Stewart and Neil Burgess for their valuable comments which helped to improve the paper. We also thank each of the data providers for providing their data openly for further use. Open access funds were provided by GEOEssential H2020 ERA-PLANET Project Number 689443. ADR acknowledges the support of REV Ocean during this work.

## Author contributions

J.A.K. and F.F. contributed equally: conceptualization, data processing, methodology, data interpretation and analysis, validation, writing - original draft preparation, figures.; J.A.K. coordinated the work; S.P. contributed to the methodology, validation, and writing - review and editing; A.D.R. contributed to the conceptualization, methodology, and writing - review and editing; L.V.W contributed to the methodology and writing - review and editing; O.A.-O. contributed to the conceptualization and writing - review and editing; A.N. contributed to the methodology, data interpretation and analysis, writing - review and editing, figures, and supervision.

