## supplementary information 1 for "Habitat Protection Indexes - new monitoring measures for the conservation of coastal and marine habitats"

### Data Sources and References

a. **SI Table 1:** Description of all data sources used in the workflow.

| Name | Website and Reference | Date of Access | Version |
|---|---|---|---|
| World Database on Protected Areas | Available at: www.protectedplanet.net[1] | May2021 | May 2021 |
| World Database on Other Effective Area-Based Conservation Measures | Available at: https://www.protectedplanet.net/en/thematic-areas/oecms?tab=OECMs[2] | May 2021 | May 2021 |
| Union of EEZs and countries | Available online at: https://www.marineregions.org/[3] | December 2020 | Version 3 |
| EEZs | Available online at: https://www.marineregions.org/[4] | March 2021 | Version 11 |
| Cold Corals | Available online at: https://doi.org/10.34892/72x9-rt61[5] | December 2020 | Version 5 |
| Warm-water Corals | Available online at: https://doi.org/10.34892/t2wk-5t34[6] | December 2020 | Version 4 |
| Knolls and Seamounts | Available online at: https://data.unep-wcmc.org/datasets/41[7] | March 2021 | Version 1.0 |

| | | | |
|---|---|---|---|
| Mangroves | Available online at: https://data.unep-wcmc.org/datasets/45[8] | December 2020 | GMW 2016 |
| Saltmarshes | Available online at: https://doi.org/10.34892/07vk-ws51[9] | December 2020 | Version 6 |
| Seagrasses | Available online at: https://doi.org/10.34892/x6r3-d211[10] | December 2020 | Version 7 |
| Ocean | Available online at: https://www.naturalearthdata.com/downloads/110m-physical-vectors/110m-ocean/[11] | December 2020 | Version 4.1.0 |
