## supplementary information 2 for "Habitat Protection Indexes - new monitoring measures for the conservation of coastal and marine habitats"

### Global Habitat Protection Index (GHPI) and Local Habitat Protection Index (LHPI) per Habitat

In this supplementary material, we present the Global Habitat Protection Index (GHPI) and Local Habitat Protection Index (LHPI) for each habitat considered in the analysis.

For quick reference the GHPI indicates the amount of habitat within PAs or OECMs divided by the total global area of the same habitat and the LHPI indicates the amount of habitat within PAs or OECMs divided by the area of the habitat within each jurisdiction.

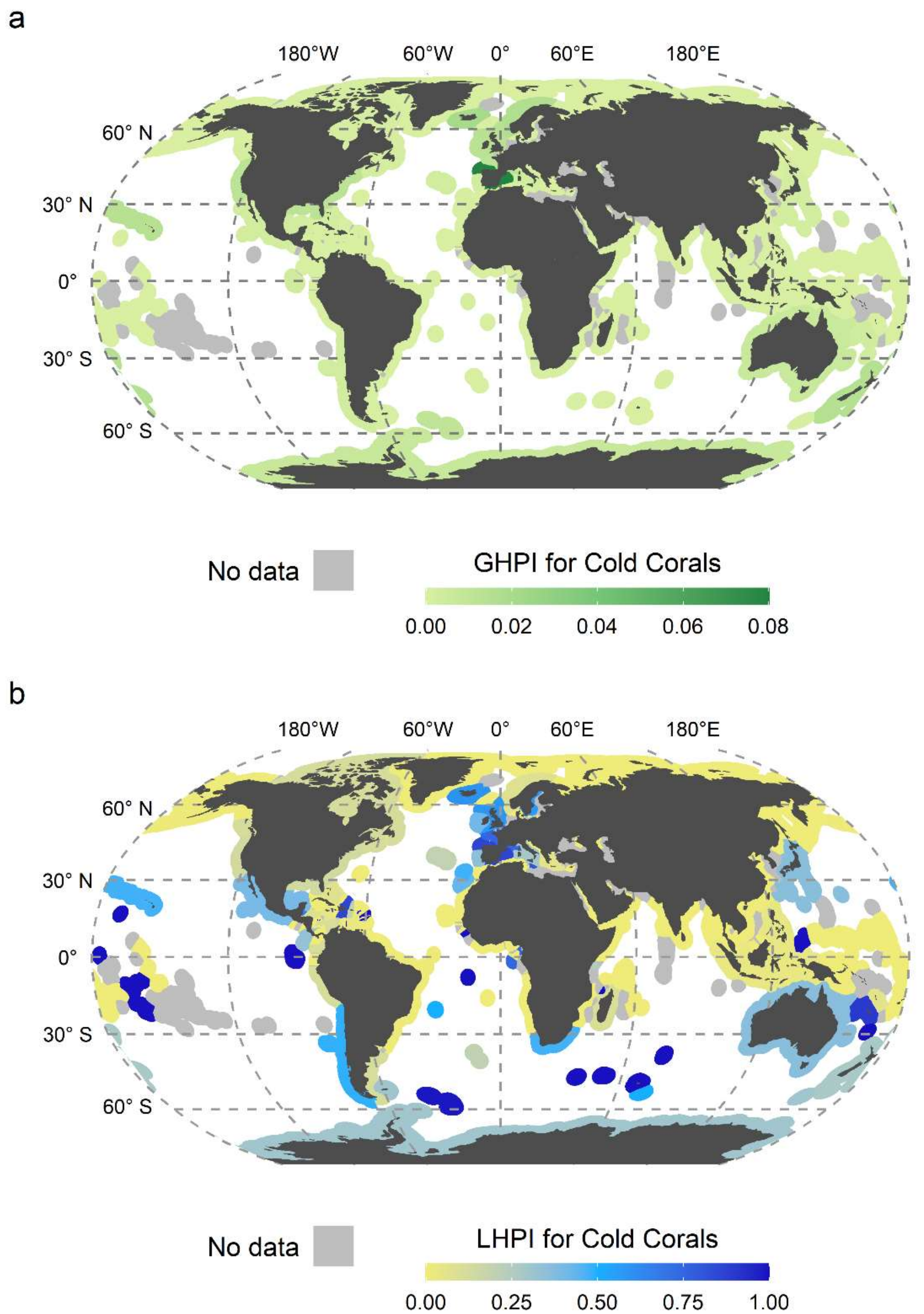

a. **Supplementary Figure 1:** Global habitat protection index and local habitat protection index for cold corals. a) GHPI illustrates the contribution of jurisdictions to the global protection of cold corals, ranging from yellow-green (low contribution) to dark green (high contribution). The index ranges from 0 to 1, but only 0 to ~0.08 is depicted here due to no jurisdictions scoring higher than ~0.08. b) LHPI illustrates how much a jurisdiction is covering their cold corals with PAs or OECMs compared to the maximum habitat extent, ranging from yellow (low contribution) to dark blue (high) contribution.

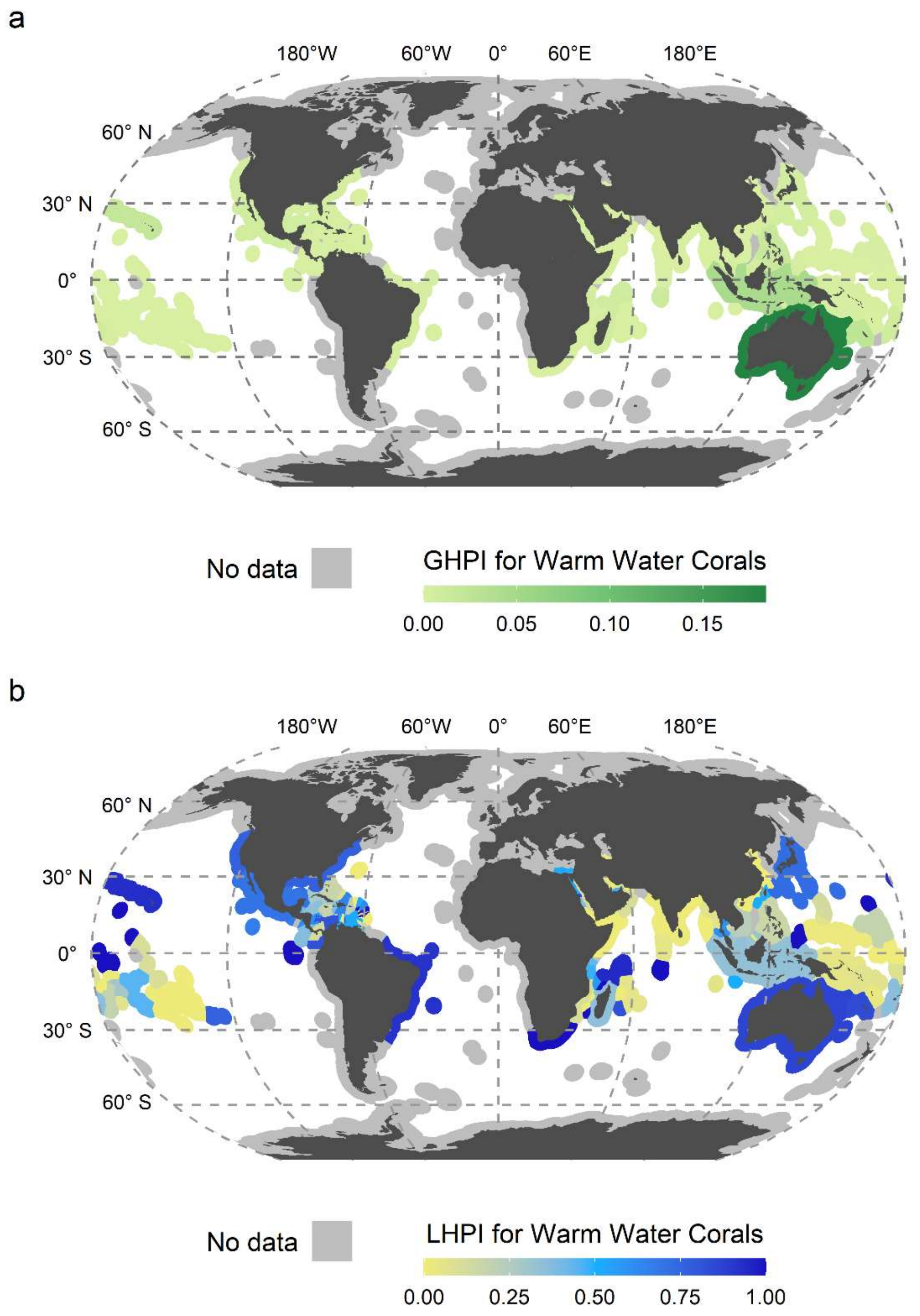

b. **Supplementary Figure 2:** Global habitat protection index and local habitat protection index for warm water corals. a) GHPI illustrates the contribution of jurisdictions to the global protection of warm water corals, ranging from yellow-green (low contribution) to dark green (high contribution). The index ranges from 0 to 1, but only 0 to ~0.18 is depicted here due to no jurisdictions scoring higher than ~0.18. b) LHPI illustrates how much a jurisdiction is covering their warm water corals with PAs or OECMs compared to the maximum habitat extent, ranging from yellow (low contribution) to dark blue (high) contribution.

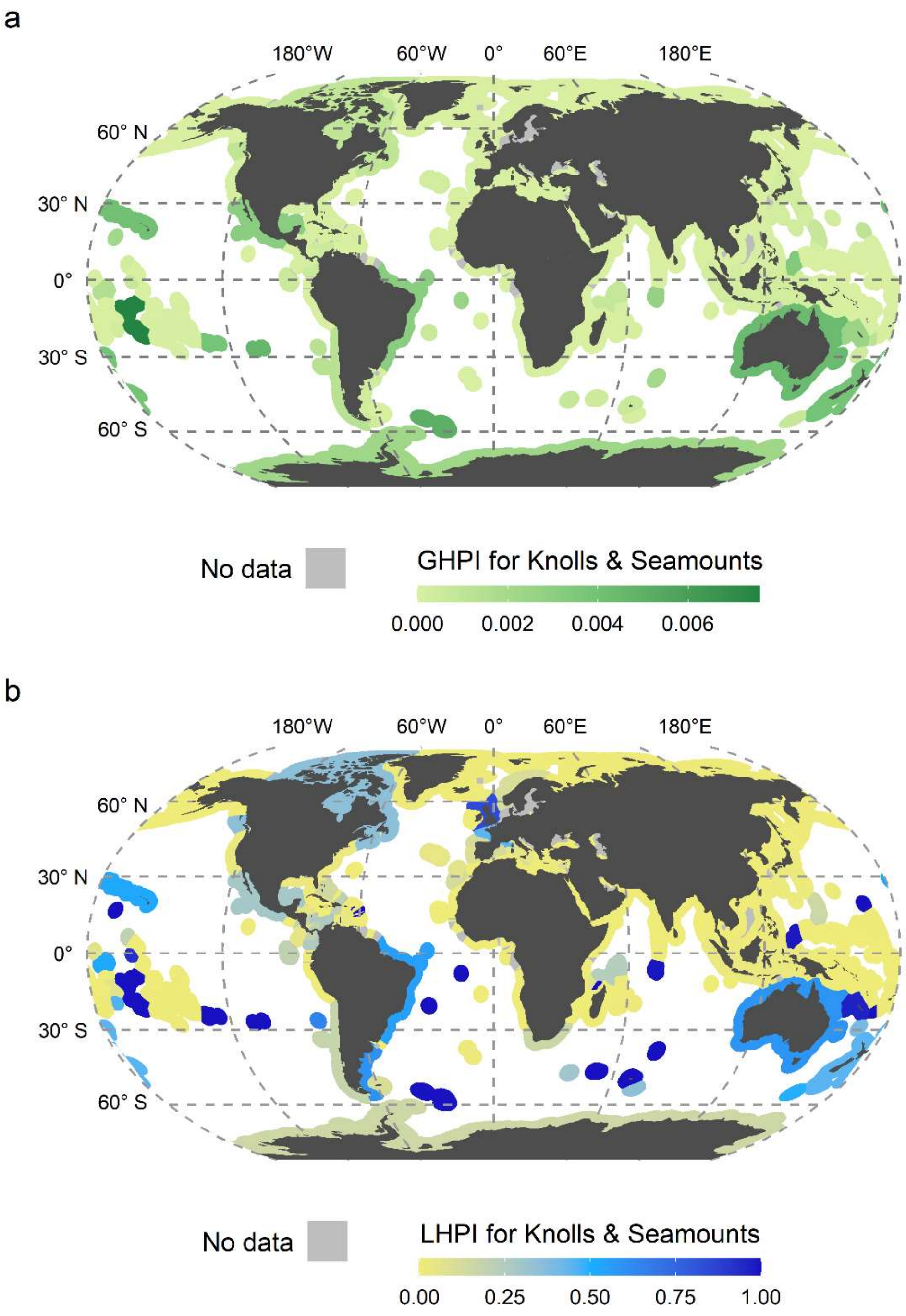

c. **Supplementary Figure 3:** Global habitat protection index and local habitat protection index for knolls and seamounts. a) GHPI illustrates the contribution of jurisdictions to the global protection of knolls and seamounts, ranging from yellow-green (low contribution) to dark green (high contribution). The index ranges from 0 to 1, but only 0 to ~0.008 is depicted here due to no jurisdictions scoring higher than ~0.008. b) LHPI illustrates how much a jurisdiction is covering their knolls and seamounts with PAs or OECMs compared to the maximum habitat extent, ranging from yellow (low contribution) to dark blue (high) contribution.

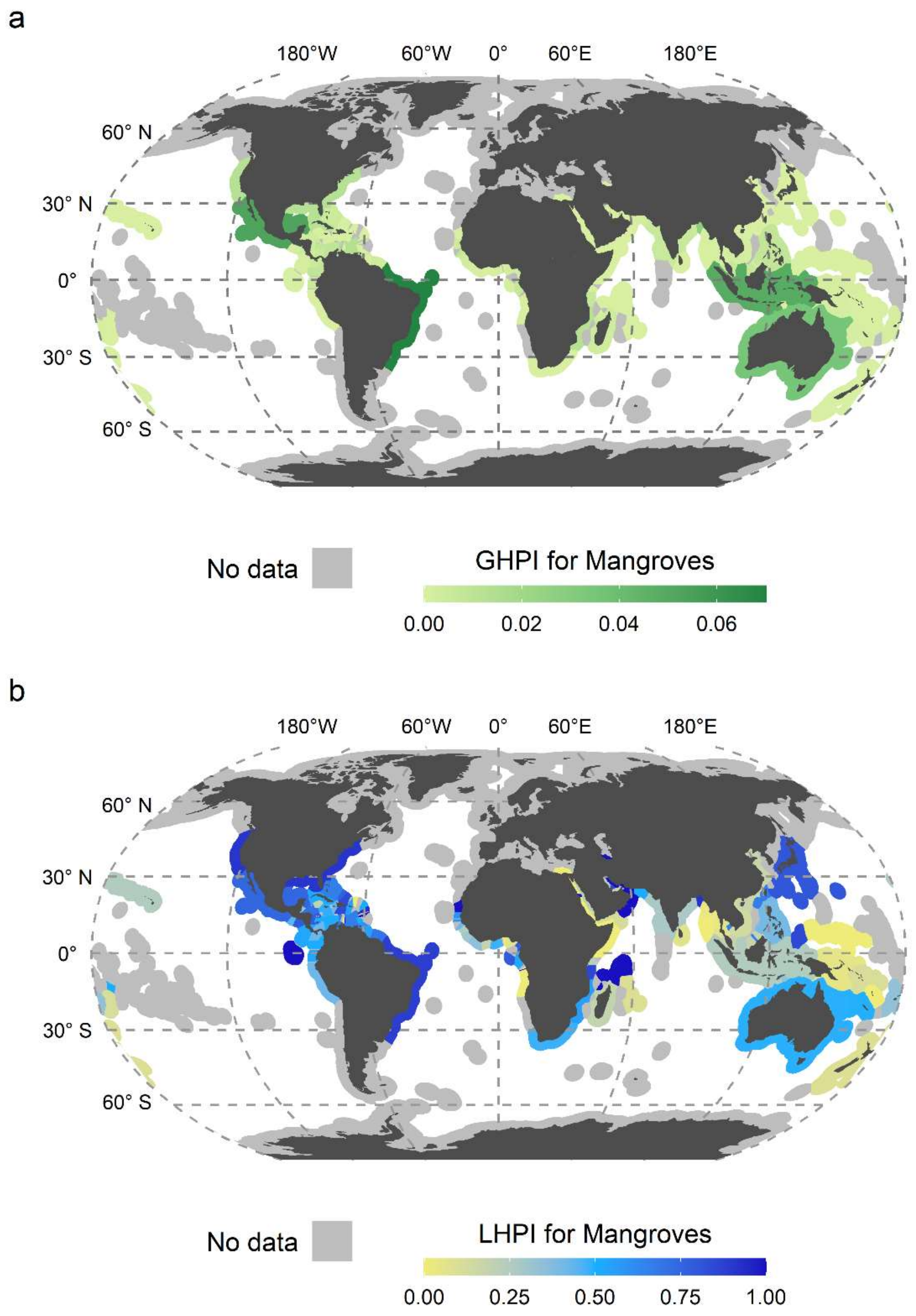

d. **Supplementary Figure 4:** Global habitat protection index and local habitat protection index for mangroves. a) GHPI illustrates the contribution of jurisdictions to the global protection of mangroves, ranging from yellow-green (low contribution) to dark green (high contribution). The index ranges from 0 to 1, but only 0 to ~0.07 is depicted here due to no jurisdictions scoring higher than ~0.07. b) LHPI illustrates how much a jurisdiction is covering their mangroves with PAs or OECMs compared to the maximum habitat extent, ranging from yellow (low contribution) to dark blue (high) contribution.

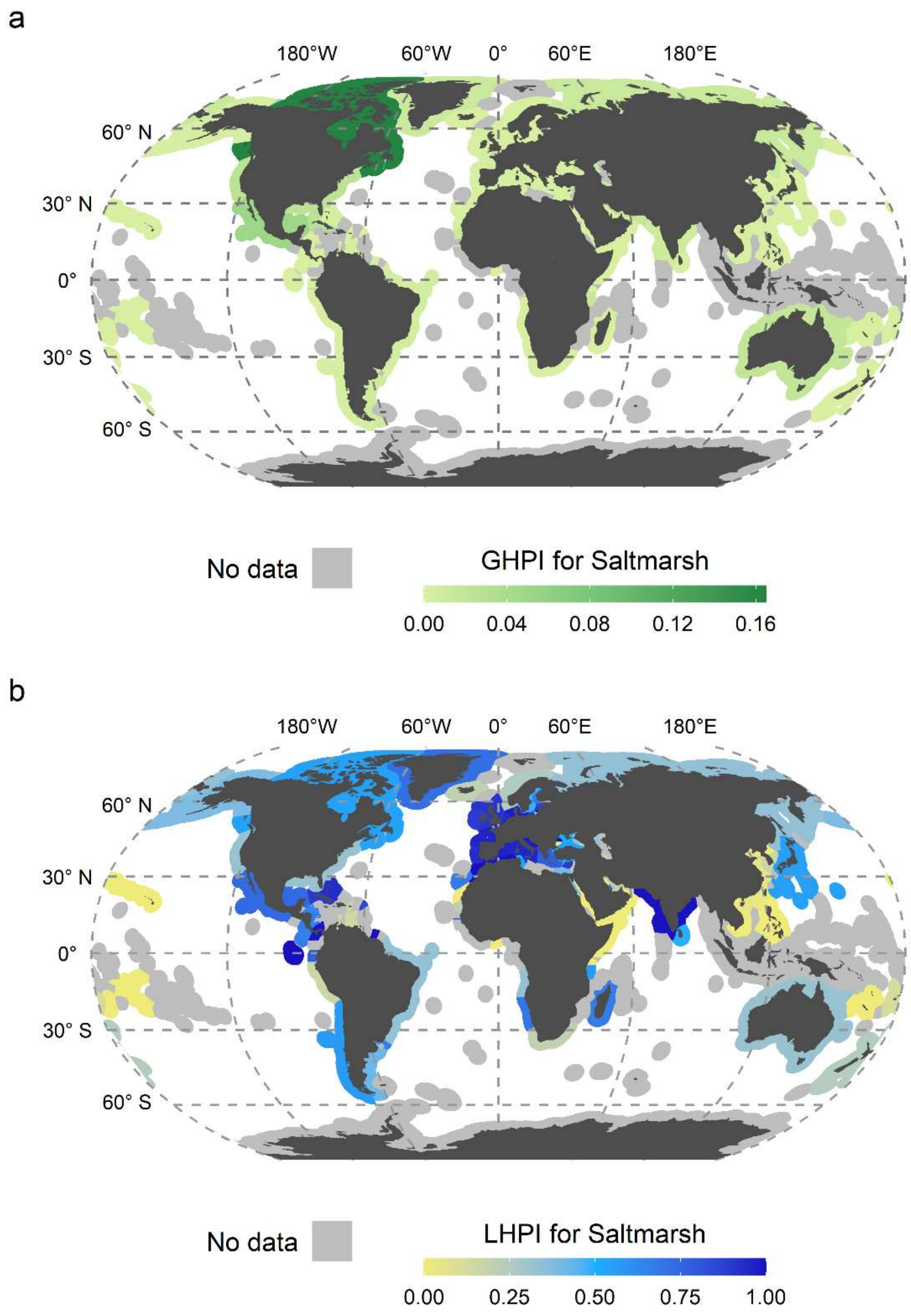

e. **Supplementary Figure 5:** Global habitat protection index and local habitat protection index for saltmarshes. a) GHPI illustrates the contribution of jurisdictions to the global protection of saltmarshes, ranging from yellow-green (low contribution) to dark-green (high contribution). The index ranges from 0 to 1, but only 0 to ~0.16 is depicted here due to no jurisdictions scoring higher than ~0.16. b) LHPI illustrates how much a jurisdiction is covering their saltmarshes with PAs or OECMs compared to the maximum habitat extent, ranging from yellow (low contribution) to dark blue (high) contribution.

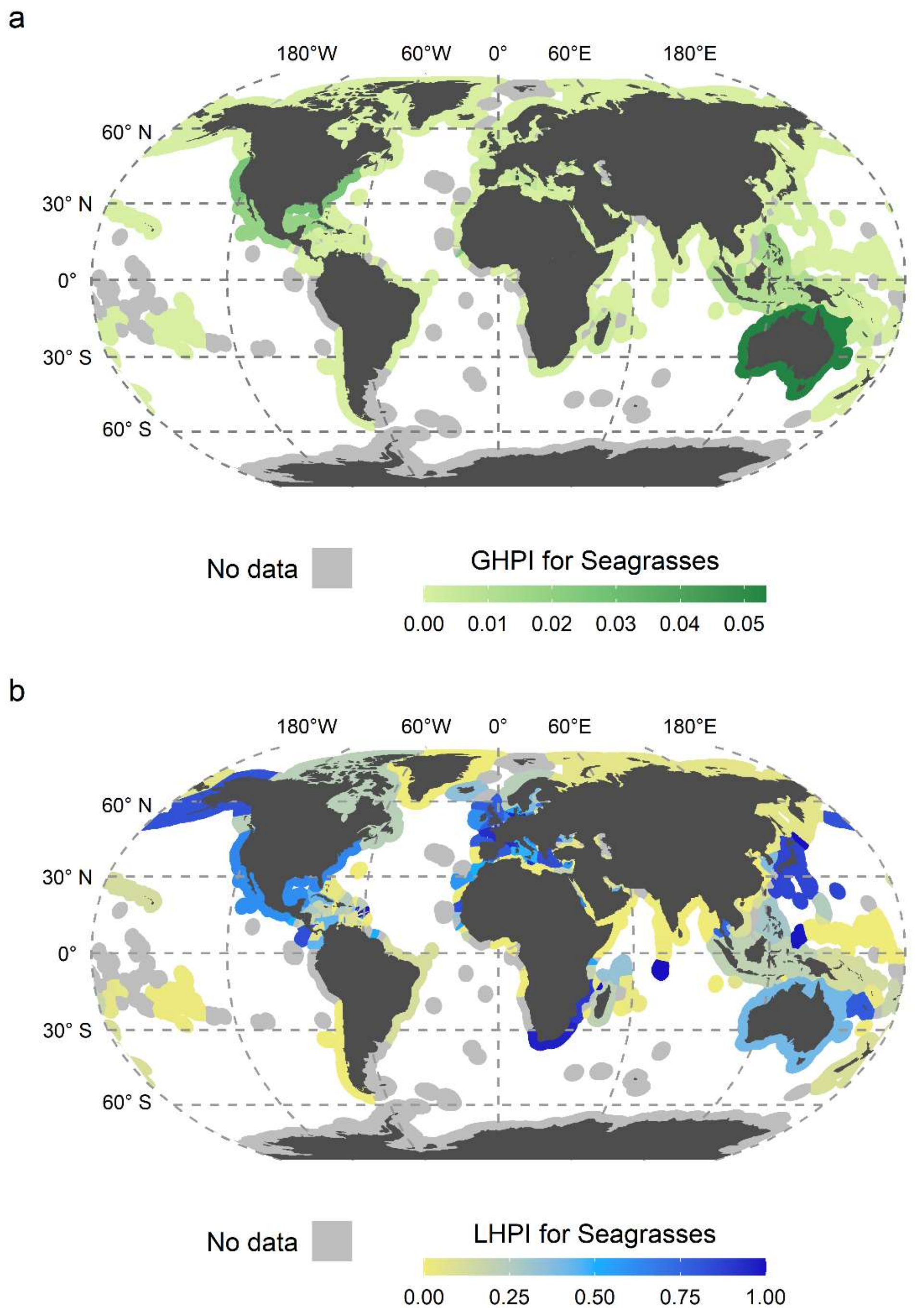

f. **Supplementary Figure 6:** Global habitat protection index and local habitat protection index for seagrasses. a) GHPI illustrates the contribution of jurisdictions to the global protection of seagrasses, ranging from yellow-green (low contribution) to dark green (high contribution). The index ranges from 0 to 1, but only 0 to ~ 0.05 is depicted here due to no jurisdictions scoring higher than ~0.05. b) LHPI illustrates how much a jurisdiction is covering their seagrasses with PAs or OECMs compared to the maximum habitat extent, ranging from yellow (low contribution) to dark blue (high) contribution.
