## supplementary information 3 for "Habitat Protection Indexes - new monitoring measures for the conservation of coastal and marine habitats"

### 30% Protection Target Figures per Habitat

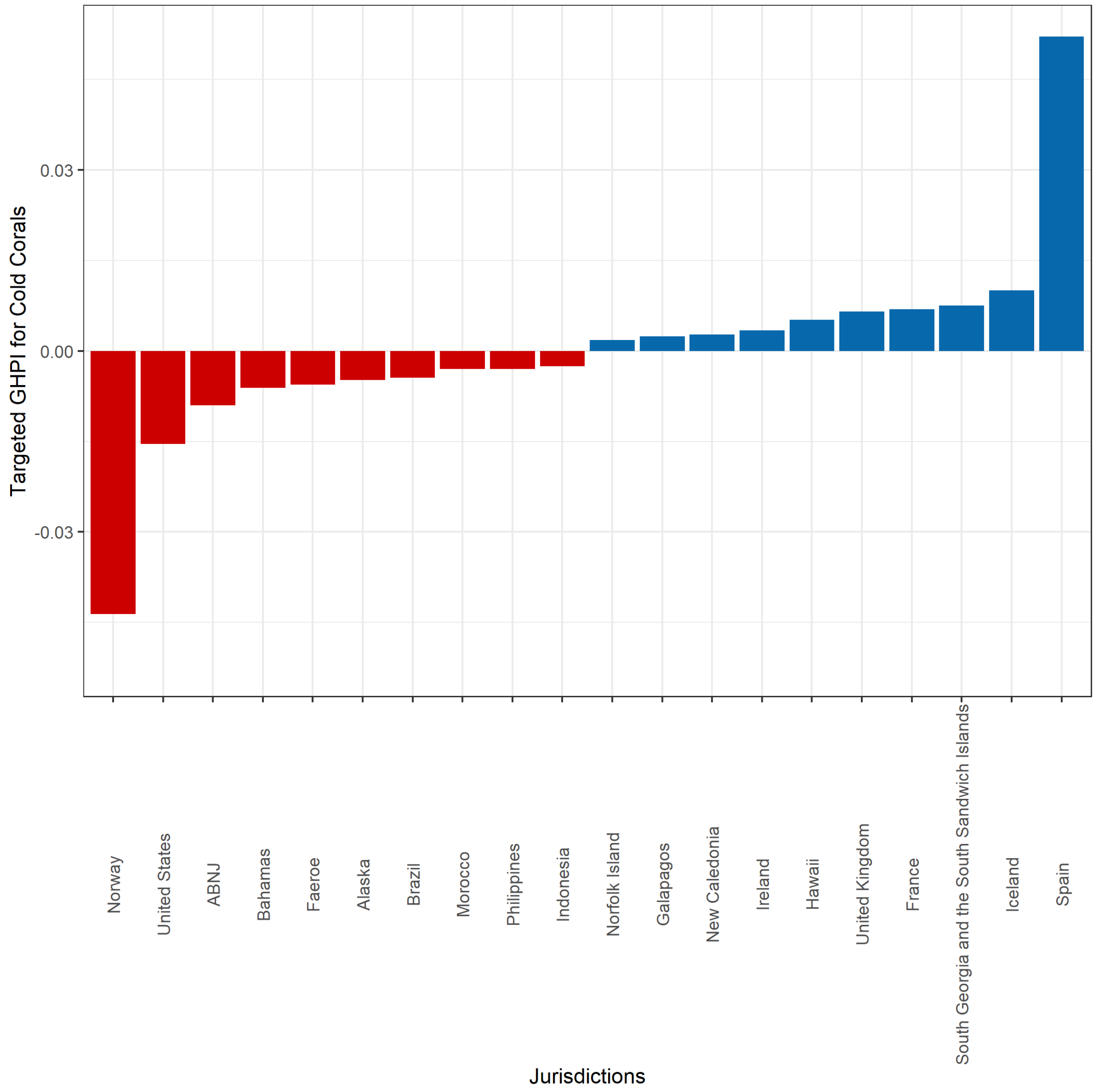

a. **Supplementary Figure 7:** Targeted global habitat protection for cold corals. The top 10 and bottom 10 out of 156 jurisdictions ranked according to their targeted Global Habitat Protection Index illustrating whether these countries have on average 30% of their cold corals within PAs or OECMs. Jurisdictions with a positive THPI indicate (blue) that more than 30% of their cold coral extent fall within PAs or OECMs, while jurisdictions with a negative THPI indicate that less than 30% of their cold coral extent fall within PAs or OECMs.

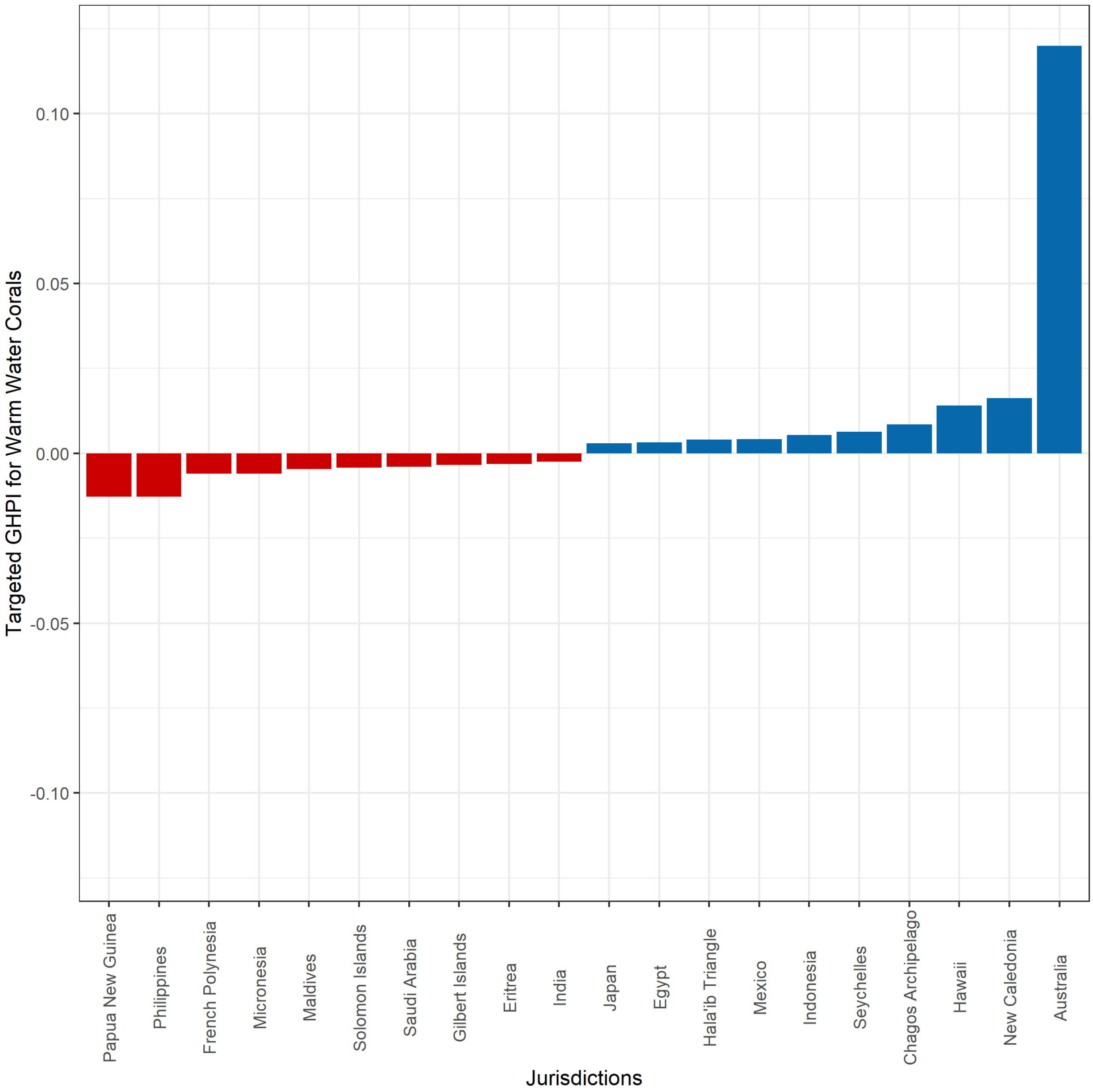

b. **Supplementary Figure 8:** Targeted global habitat protection for warm water corals. The top 10 and bottom 10 out of 134 jurisdictions ranked according to their targeted Global Habitat Protection Index illustrating whether these countries have on average 30% of their warm water corals within PAs or OECMs. Jurisdictions with a positive THPI indicate (blue) that more than 30% of their warm water coral extent fall within PAs or OECMs, while jurisdictions with a negative THPI indicate that less than 30% of their warm water coral extent fall within PAs or OECMs.

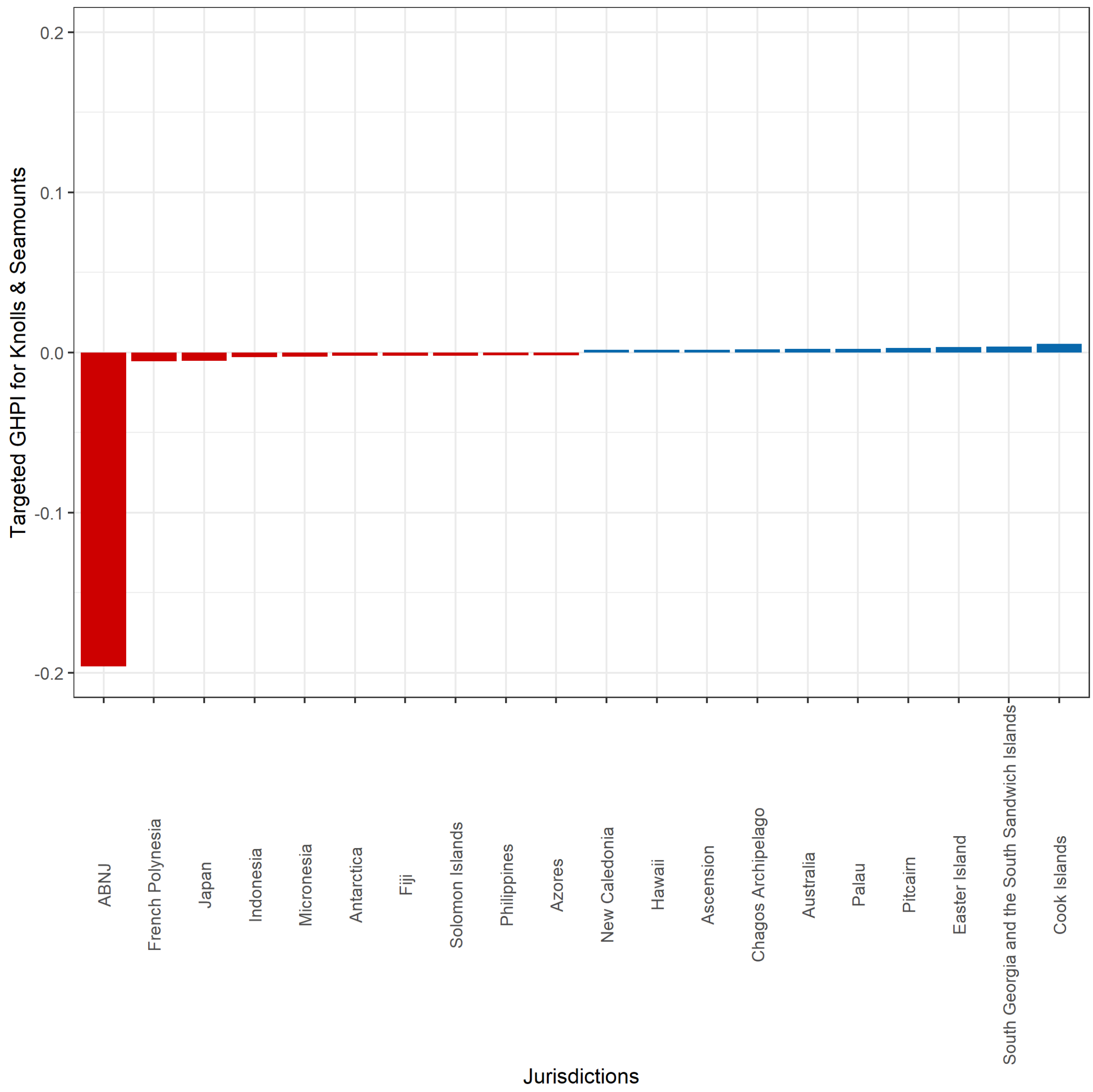

c. **Supplementary Figure 9:** Targeted global habitat protection for knolls and seamounts. The top 10 and bottom 10 out of 196 jurisdictions ranked according to their targeted Global Habitat Protection Index illustrating whether these countries have on average 30% of their knolls and seamounts within PAs or OECMs. Jurisdictions with a positive THPI indicate (blue) that more than 30% of their knoll and seamount extent fall within PAs or OECMs, while jurisdictions with a negative THPI indicate that less than 30% of their knoll and seamount extent fall within PAs or OECMs.

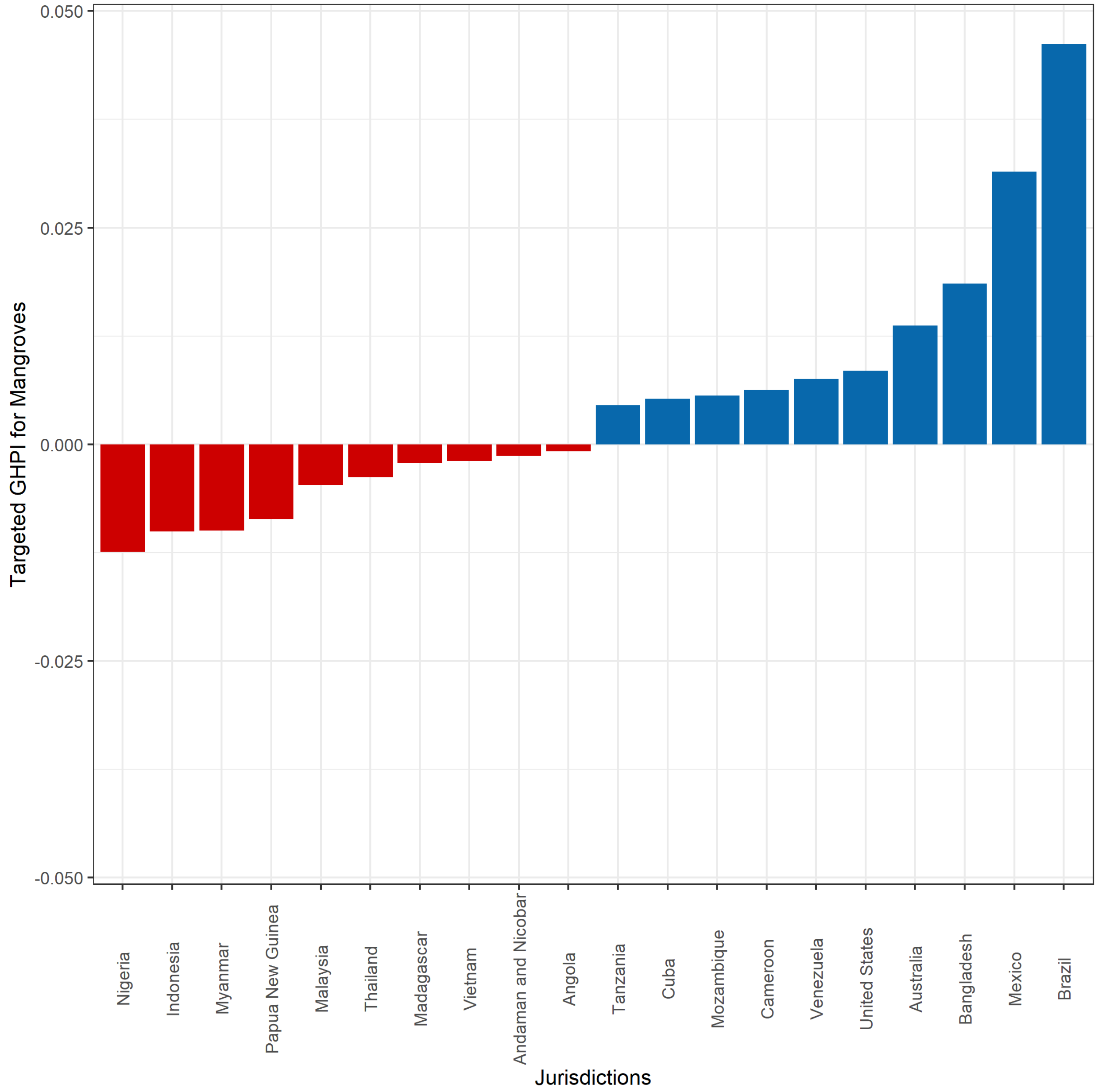

d. **Supplementary Figure 10:** Targeted global habitat protection for mangroves. The top 10 and bottom 10 out of 103 jurisdictions ranked according to their targeted Global Habitat Protection Index illustrating whether these countries have on average 30% of their mangroves within PAs or OECMs. Jurisdictions with a positive THPI indicate (blue) that more than 30% of their mangrove extent fall within PAs or OECMs, while jurisdictions with a negative THPI indicate that less than 30% of their mangrove extent fall within PAs or OECMs.

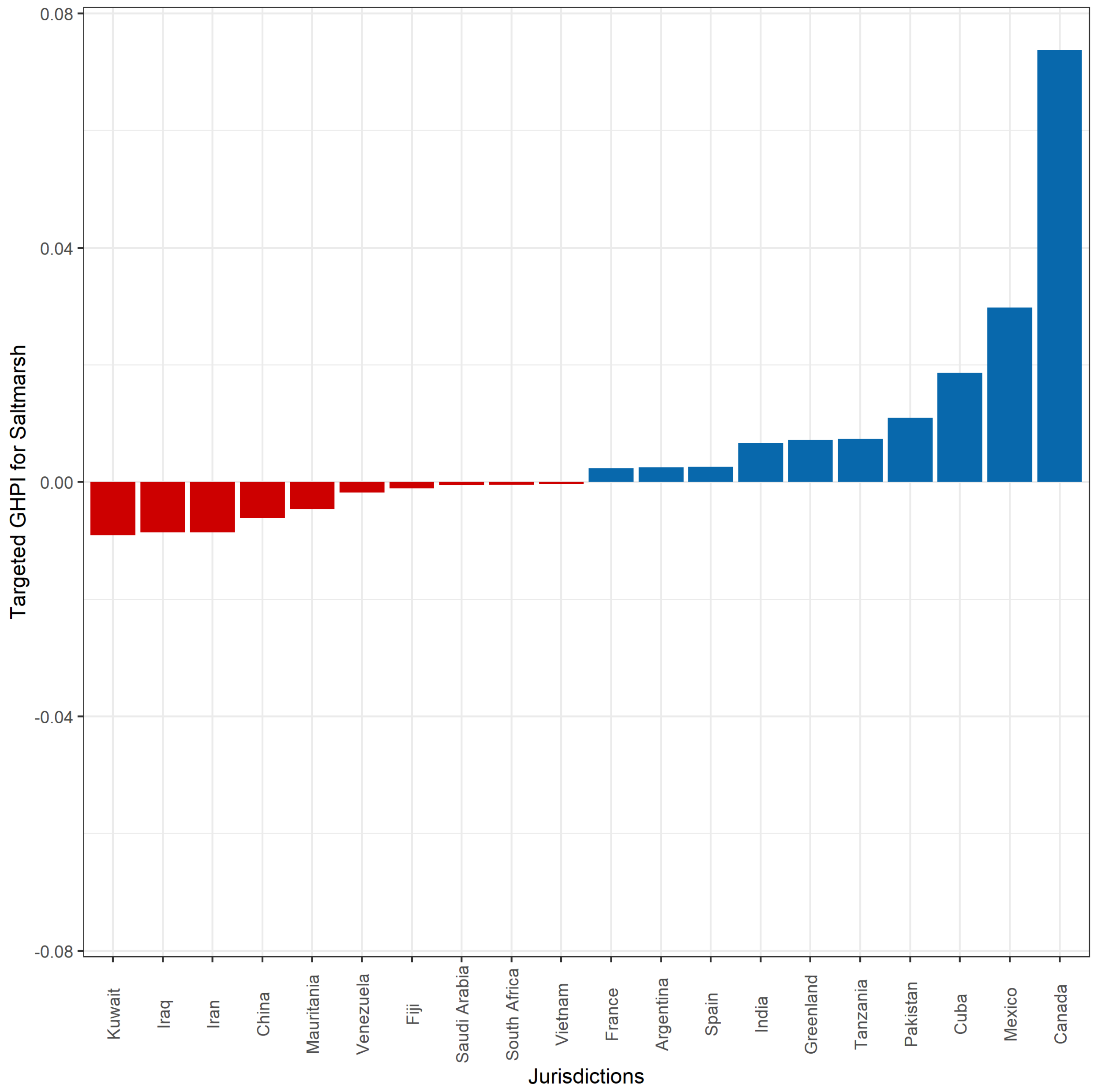

e. **Supplementary Figure 11:** Targeted global habitat protection for saltmarsh. The top 10 and bottom 10 out of 99 jurisdictions ranked according to their targeted Global Habitat Protection Index illustrating whether these countries have on average 30% of their saltmarsh within PAs or OECMs. Jurisdictions with a positive THPI indicate (blue) that more than 30% of their saltmarsh extent fall within PAs or OECMs, while jurisdictions with a negative THPI indicate that less than 30% of their saltmarsh extent fall within PAs or OECMs.

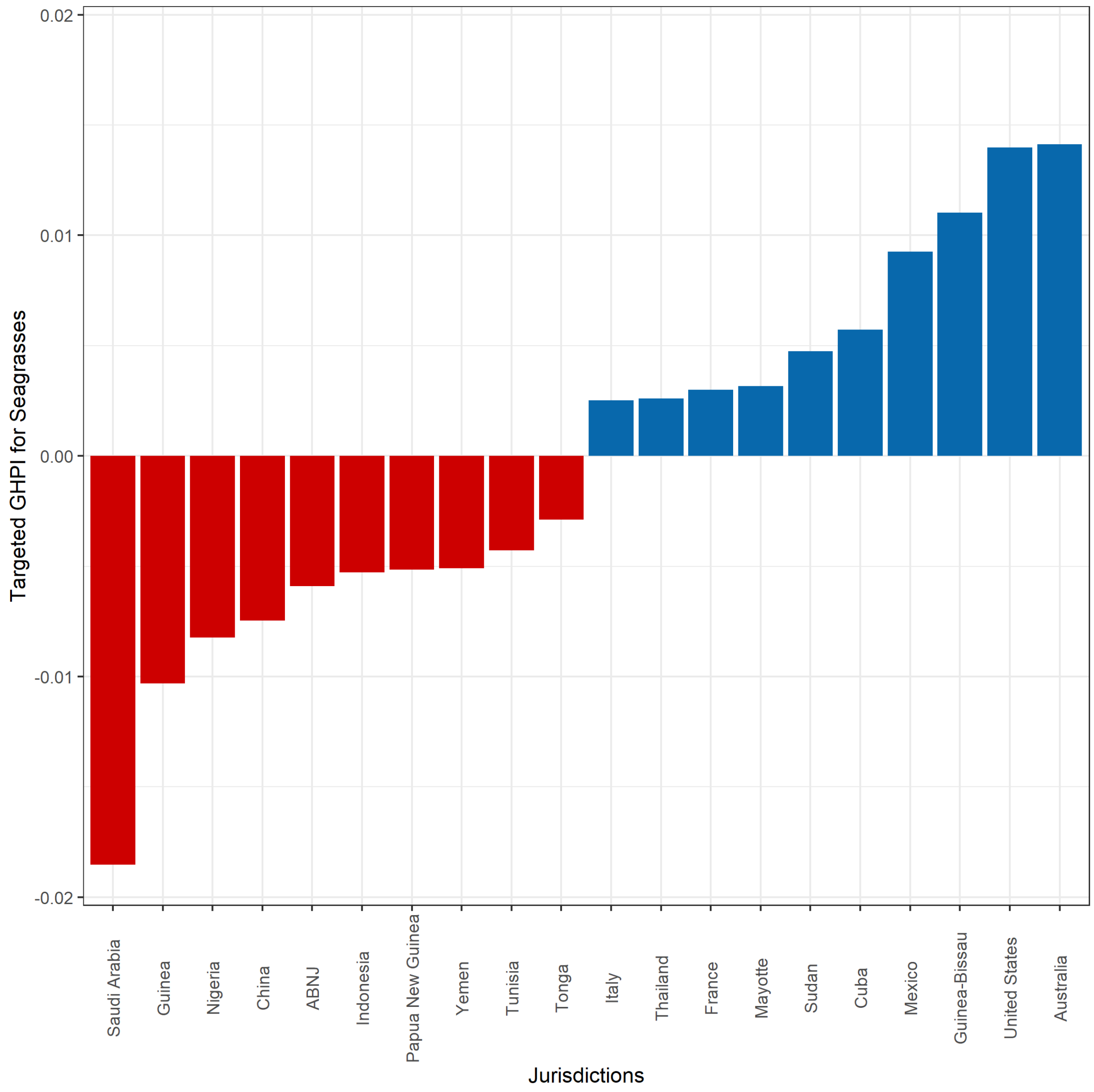

f. **Supplementary Figure 12:** Targeted global habitat protection for seagrasses. The top 10 and bottom 10 out of 165 jurisdictions ranked according to their targeted Global Habitat Protection Index illustrating whether these countries have on average 30% of their seagrasses within PAs or OECMs. Jurisdictions with a positive THPI indicate (blue) that more than 30% of their seagrass extent fall within PAs or OECMs, while jurisdictions with a negative THPI indicate that less than 30% of their seagrass extent fall within PAs or OECMs.
